# Arp2/3 mediated dynamic lamellipodia of the hPSC colony edges promote liposome-based DNA delivery

**DOI:** 10.1101/2021.05.16.444342

**Authors:** Michelle Surma, Kavitha Anbarasu, Arupratan Das

**Author notes:** Michelle Surma: Collection and/or assembly of data, data analysis and interpretation, administrative support, manuscript writing. Kavitha Anbarasu: Collection and/or assembly of data. Arupratan Das: Conception and design, financial support, administrative support, collection and/or assembly data, data analysis and interpretation, manuscript writing, and final approval of manuscript. Correspondence: Arupratan Das, PhD. Indiana University School of Medicine 1160 W. Michigan Street, GK305W, Indianapolis, IN 46202. This work was supported by grant from the NIH, United States (R00EY028223) and startup package to AD from the Indiana University School of Medicine.

## Abstract

Cationic liposome-mediated delivery of drugs, DNA, or RNA plays a pivotal role in small molecule therapy, gene editing, and immunization. However, our current knowledge regarding the cellular structures that facilitate this process remains limited. Here, we used human pluripotent stem cells (hPSCs), which form compact colonies consisting of dynamically active cells at the periphery and epithelial-like cells at the core. We discovered that cells at the colony edges selectively got transfected by cationic liposomes through Arp2/3 dependent dynamic lamellipodia, which is augmented by myosin II inhibition. Conversely, cells at the core establish tight junctions at their apical surfaces, impeding liposomal access to the basal lamellipodia and thereby inhibiting transfection. In contrast, liposomes incorporating mannosylated lipids are internalized throughout the entire colony via receptor-mediated endocytosis. These findings contribute a novel mechanistic insight into enhancing therapeutic delivery via liposomes, particularly in cell types characterized by dynamic lamellipodia, such as immune cells, or those comprising the epithelial layer.

**Significance Statement:** Drug or gene delivery to human cells is essential for effective treatment. Cationic liposomes provide a safe delivery vehicle compared to viruses. However, the cellular structures required for internalizing liposomes are not yet fully understood. Using human stem cells which grow in compact colonies with more dynamic cells at the periphery and epithelial like cells at the center, here we discovered that Arp2/3 dependent dynamic lamellipodia promotes cationic liposome delivery in dynamic cells while receptor mediated endocytosis is required for epithelial cells. This is significant as it provides mechanisms for enhancing liposome delivery to both migratory and epithelial cells in our body.

Graphical AbstractMechanisms for liposome transfection to the lamellipodial or epithelial cells.
Data shown here suggest cationic liposomes fuse with the negatively charged dynamic lamellipodia membrane in an Arp2/3 dependent manner and the process is enhanced by Myosin II inhibition, such as with the stem cell colony edge cells. However, cells more epithelial in nature such as those inside the stem cell colony center do not possess dynamic lamellipodia at the apical surface, rather they form tight junctions which inhibit cationic liposome transfection. Epithelial cells rely on receptor mediated endocytosis in both Myosin II dependent and independent manners to internalize liposomes with lipids that contain ligands for cell surface receptors such as mannose.

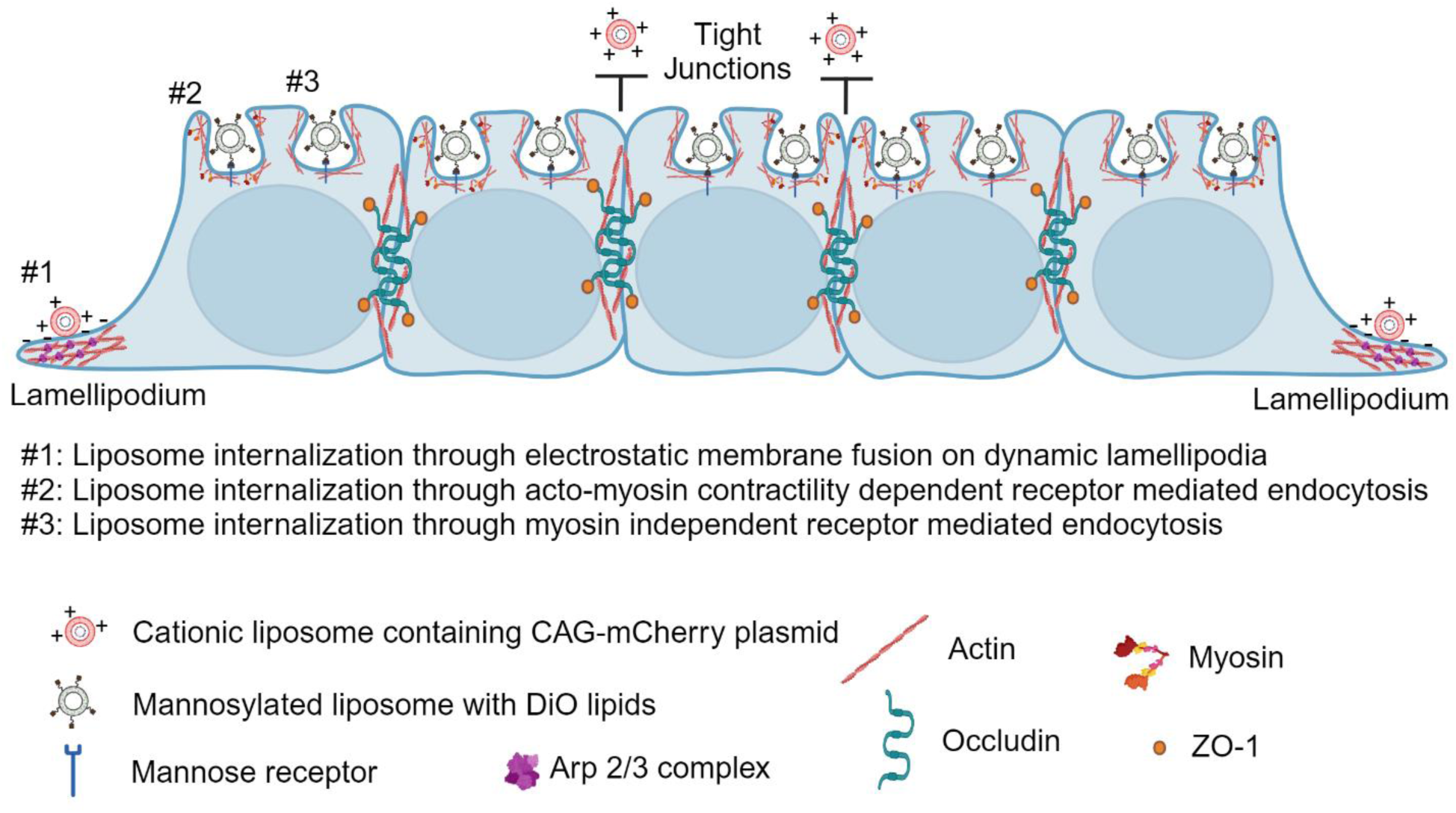

## Introduction

Liposomes serve as highly efficient and safe delivery vehicles for a wide spectrum of therapeutics, encompassing drugs and genetic materials such as genes and messenger RNAs (mRNAs). A compelling example lies in the development of anti-COVID-19 vaccines, which have played a pivotal role in combating the ongoing pandemic crisis. In this effort, mRNA encoding the spike protein of the Coronavirus is encapsulated within cationic liposomes, facilitating their delivery into the bloodstream. Subsequently, these liposomes are internalized by immune cells, triggering the production of antibodies against the virus. Notably, cationic lipids significantly augment the fusion of these liposomes with the anionic surface of the plasma membrane lipid layer, culminating in the liberation of their payload into the cytosol. Moreover, cationic liposomes have the capacity to enter cells through endocytosis while escaping lysosomal degradation. When cationic liposomes are endocytosed, their uptake by cells leads to an intriguing series of events. The endosomal membrane, originating from the plasma membrane, interacts with the cationic liposomal lipids, inducing membrane destabilization. Consequently, a flip-flop phenomenon occurs, wherein the anionic lipids from the cytosolic surface of the endosomal membrane reorient to face its inner surface, while the liposomal cationic lipids project outward into the endosomal membrane. This interaction triggers the lateral diffusion of anionic lipids (from the endosome membrane) and cationic lipids (from the liposome), resulting in the formation of charge-neutralized lipid pairs. This, in turn, leads to the disruption of the lipid bilayer, facilitating the release of liposomal content into the cytoplasm. These intricate mechanisms allow liposomes to circumvent endo-lysosomal degradation, ensuring the efficient release of cargo into the cytoplasm (Zelphati and Szoka, 1996). While substantial progress has been made in elucidating the compositions and delivery mechanisms of liposomes via endocytosis, there remains a lack of comprehensive knowledge concerning the cell surface structures that are indispensable for cationic liposome internalization.

Numerous cell-surface mechanisms may contribute to the internalization of cationic liposomes. These mechanisms include fusion with the anionic plasma membrane surface (Bennett et al., 1992; Felgner and Ringold, 1989), endocytosis (Zabner et al., 1995; Zelphati and Szoka, 1996) or receptor-mediated endocytosis when liposomes contain modified lipids (Kelly et al., 2011). These processes can occur either at the apical surface of epithelial cell layers or at membrane structures found at the leading edge of migratory cells, such as lamellipodia or filopodia. It is noteworthy that endocytic internalization is contingent upon actomyosin contractility (Doherty and McMahon, 2009), while the fusion of liposomes with the plasma membrane can occur independently of actomyosin processes (Felgner and Ringold, 1989). The therapeutic delivery of nucleotides including DNA and mRNA, or drugs, frequently employs cationic liposomes as carriers (Sercombe et al., 2015). An in-depth comprehension of the cellular structures and associated molecular mechanisms necessary for the internalization of cationic liposomes holds significant promise for the advancement of liposome-based therapeutics. These potential benefits encompass the reduction of drug dosages, the enhancement of immunization efficacy, and the augmentation of CRISPR-based gene-editing efficiency. This knowledge has far-reaching implications for gene therapy and disease modeling research.

In this study, we harnessed the unique properties of human pluripotent stem cells (hPSCs), which form compact colonies characterized by the presence of epithelial cell junctions at the colony center, and more dynamic cells at the periphery (Kim et al., 2022). Our investigation revealed a distinctive pattern: cells situated at the colony periphery, as opposed to those at the center, exhibited dynamic lamellipodia structures and demonstrated selective transfection by cationic liposomes. Remarkably, this behavior was found to be independent of actomyosin contractility. Instead, it was notably amplified under myosin II inhibition, and critically reliant on the Arp2/3-mediated dynamic actin meshwork-driven lamellipodial structures. This edge-specific transfection phenomenon was unique to cationic liposomes. However, in contrast to this selective transfection, we observed that cells distributed throughout the colony were efficiently transfected by liposomes containing mannosylated lipids, achieved through receptor-mediated endocytosis. Given that endocytosis is typically dependent on actomyosin contractility, our findings suggesting enhanced cationic liposome delivery to hPSCs under myosin II inhibition point towards an intriguing actomyosin-independent, passive membrane fusion mechanism. It’s noteworthy that although hPSCs across the entire colony exhibited transfection with mannosylated liposomes, this process was somewhat influenced by myosin II inhibition, albeit not entirely. This observation implies the presence of both actomyosin contractility-dependent and -independent receptor-mediated endocytic delivery mechanisms.

Thus, our study offers a unique insight into mechanisms whereby Arp2/3-mediated lamellipodial structures, akin to those found in migratory cells or the peripheral cells of epithelial tissues, can be harnessed to enhance cationic liposome-based delivery. Simultaneously, it underscores the potential for enhancing liposomal delivery in epithelial cells through receptor-mediated endocytosis, particularly via mannose receptors at the apical surface. The enhanced efficacy of liposome-based therapeutic delivery, as demonstrated in our study, holds the promise of optimizing treatments by reducing both dosage and treatment duration.

## Results

### Cells at the hPSC colony edges are selectively transfected by cationic liposomes

Human pluripotent stem cells grow in colonies with cells at the center forming tight junctions with apico-basal polarity similar to the epithelial cell layer, and with cells at the edge being more dynamic in nature (Kim et al., 2022). This gives a unique advantage to study the cellular structures essential for the cationic liposome transfection as used for therapeutic delivery. We have formed cationic liposomes by mixing cationic lipids (Lipostem, Thermo) and negatively charged plasmid DNA containing mCherry under CAG promoter following user manual. These liposomes are then added to the H7 human embryonic stem cells (H7-hESCs) grown in sparse culture by single cell accutase passaging or in small colonies by clump passaging. Next, 24h post transfection fluorescence expression was checked by imaging as shown in Figure 1 A. Much to our surprise, we observed that stem cells at the colony edges got selectively transfected but not at the colony center (Fig. 1 B), while sparse cell culture got transfected randomly with no such pattern (Fig. 1 C) as observed by the mCherry fluorescence. To measure if stem cells at the colony center are not transfected and hence not expressing mCherry, we drew a line across the colony center through the edges and measured fluorescence intensity profile along that line (Fig. 1 D). Indeed, we observed specific fluorescence intensity peaks on the line corresponding to the edges but not at the center (Fig. 1 E). This observation was further verified by measuring fluorescence intensity around the colony edges and centers, which showed significantly higher expression at the edges (Fig. 1F). The edge specific liposomal transfection is seen throughout the culture, as revealed by wider field of view using a 4x objective magnification image (Fig. S1). To test if liposome mediated edge specific transfection is a cell type specific phenomenon, we transfected induced pluripotent stem cell (EP1-iPSC) (Bhise et al., 2013) colonies with cationic liposomes containing CAG-mCherry plasmid. We observed similar edge specific transfection of these iPSC colonies (Fig. S2 A) establishing that the phenomenon is not specific to a particular stem cell line, rather a property of human stem cells. This observation is further verified by measuring fluorescence intensity profiles across the line through the colony which showed specific intensity peaks at the colony edges but not at the centers (Fig. S2 A, B). Similarly, fluorescence intensity measurements showed significantly high fluorescence at the edges compared to centers (Fig. S2 C). We occasionally observe some cells adjacent to edge cells but inside the colony also express fluorescence protein which could arise from the dividing transfected edge cells or through direct transfection. These data suggest stem cells at the colony edges possess unique structures compared to the center cells and enhancing those structures could increase the cationic liposome transfection efficiency to the edge cells.

**Figure 1:**
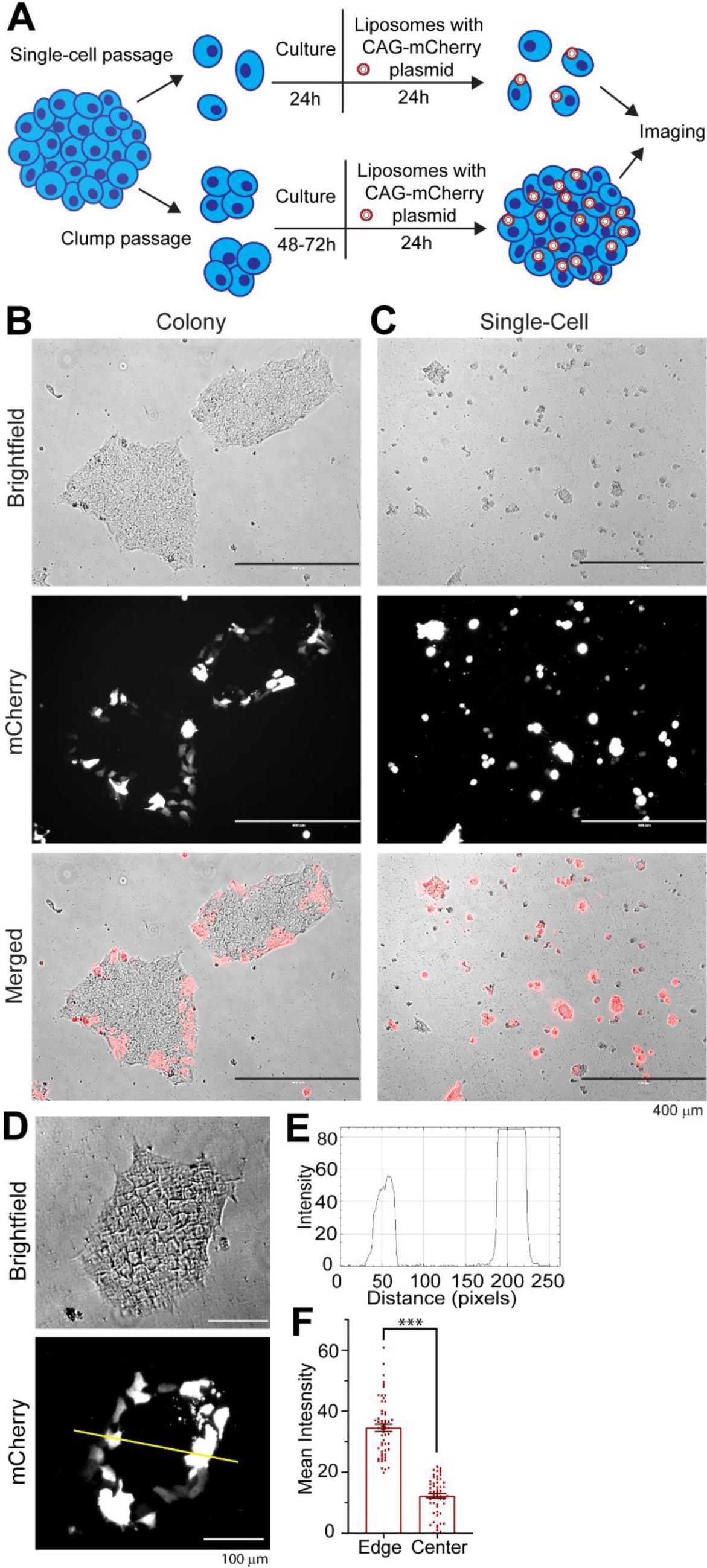
Stem cell colony edges selectively get transfected with cationic liposomes. **(A)** Illustration of stem cell transfections where H7-hESCs, after clump or single-cell passaging, were transfected with cationic liposomes containing CAG-mCherry plasmid (red fluorescence protein, RFP) plasmid and images were taken 24h after transfection. Representative 10x brightfield and RFP images of **(B)** clump-passaged colonies and **(C)** single-cell passaged stem cells. **(D)** Representative images of clump passaged colony 24h after transfection with CAG-mCherry plasmid. **(E)** Line trace through the center of colony as shown in 1D reveals fluorescence intensity peaks at the edges but not at the center. **(F)** Quantification of the fluorescence intensity at colony edges and centers from 58 colonies from 5 independent experiments. Each data point represents a single colony. Error bars are SEM. Unpaired student’s *t-test*, *** p < 0.001.

### Stem cells at the colony edges possess dynamic lamellipodia structures

It has been reported that stem cells form tight junctions at the center of the colony while cells at the edges show more dynamic behavior (Kim et al., 2022). Arp2/3 mediated polymerization of the dendritic actin network at the cell edge forms dynamic lamellipodium critical for directional cell migration (Suraneni et al., 2012). We examined whether stem cells at the colony edges form lamellipodia structures by confocal immunofluorescence imaging of actin cytoskeleton and lamellipodium marker cortactin, at the edge and center as shown in Figure 2A. Cortactin localizes to the cortical F-actin sites required for forming dynamic lamellipodia membranes (Weed and Parsons, 2001). Indeed, we observed in DMSO treated control conditions the basal surface of edge cells possess cortactin decorated actin meshwork structures at the leading edge (Fig. 2 B, white arrows), which are absent at the corresponding apical surface (Fig. 2 C). Actomyosin contractility promotes thick actin stress fiber (SF) formation (Lehtimäki et al., 2021) but reduced actomyosin contractility results in more non tensile thin actin filaments with enhanced lamellipodia structures (Goeckeler et al., 2008; Raucher and Sheetz, 2000). We asked if myosin II inhibition by a potent inhibitor blebbistatin (Kovács et al., 2004) dissolves actin SF and enriches lamellipodia at the edge cells. Indeed, we observed loss of SFs but more thin actin meshwork structures at the basal surface of edge cells (Fig. 2 B). Interestingly, we also observed more cells at the colony edges as well as immediately inside of the edges with cortactin decorated actin cortex (Fig. 2 B, white arrows) indicating more lamellipodia formation under myosin II inhibition. CK666 is a potent inhibitor of Arp2/3, and hence lamellipodia formation (Wu et al., 2012), and we saw that Arp2/3 inhibition led to more actin SF formation with reduced cortactin decorated actin cortex in the edge cells (Fig. 2 B) indicating loss of lamellipodia. We further observed myosin II inhibition by blebbistatin in presence of Arp2/3 inhibition by CK666 reduced actin SF formation to some extent, but it did not increase cortactin localization at the cell edge (Fig. 2 B). Thus, our data indicates Arp2/3 activity can promote lamellipodia in the edge cells independent of myosin II activity. The apical surface of the colony edges did not show actin SF, rather formed honeycomb like filaments characteristic of epithelial apical surface with no cortactin localization to actin (Fig. 2 C). No noticeable changes in actin or cortactin assembly were observed upon Arp2/3 and/or myosin II inhibition (Fig. 2 C). At the basal surface of colony center, cells possess extensive amount of actin SFs which did not change under Arp2/3 inhibition but were significantly reduced by myosin II inhibition alone or in combination with Arp2/3 inhibition (Fig. 2 D). However, at the basal or apical surfaces of colony center, cells did not show any cortactin localization on the actin filaments indicating the absence of lamellipodia in the center cells (Fig. 2 D, E). Interestingly, myosin II inhibition occasionally increased cortactin localization at the apical actin networks (Fig. 2 E, white arrows) indicating an increase in membrane dynamics at the apical cortex. Arp2/3 inhibition alone or in combination with myosin II inhibition did not alter actin or cortactin localizations in the apical surface (Fig. 2 E).

**Figure 2:**
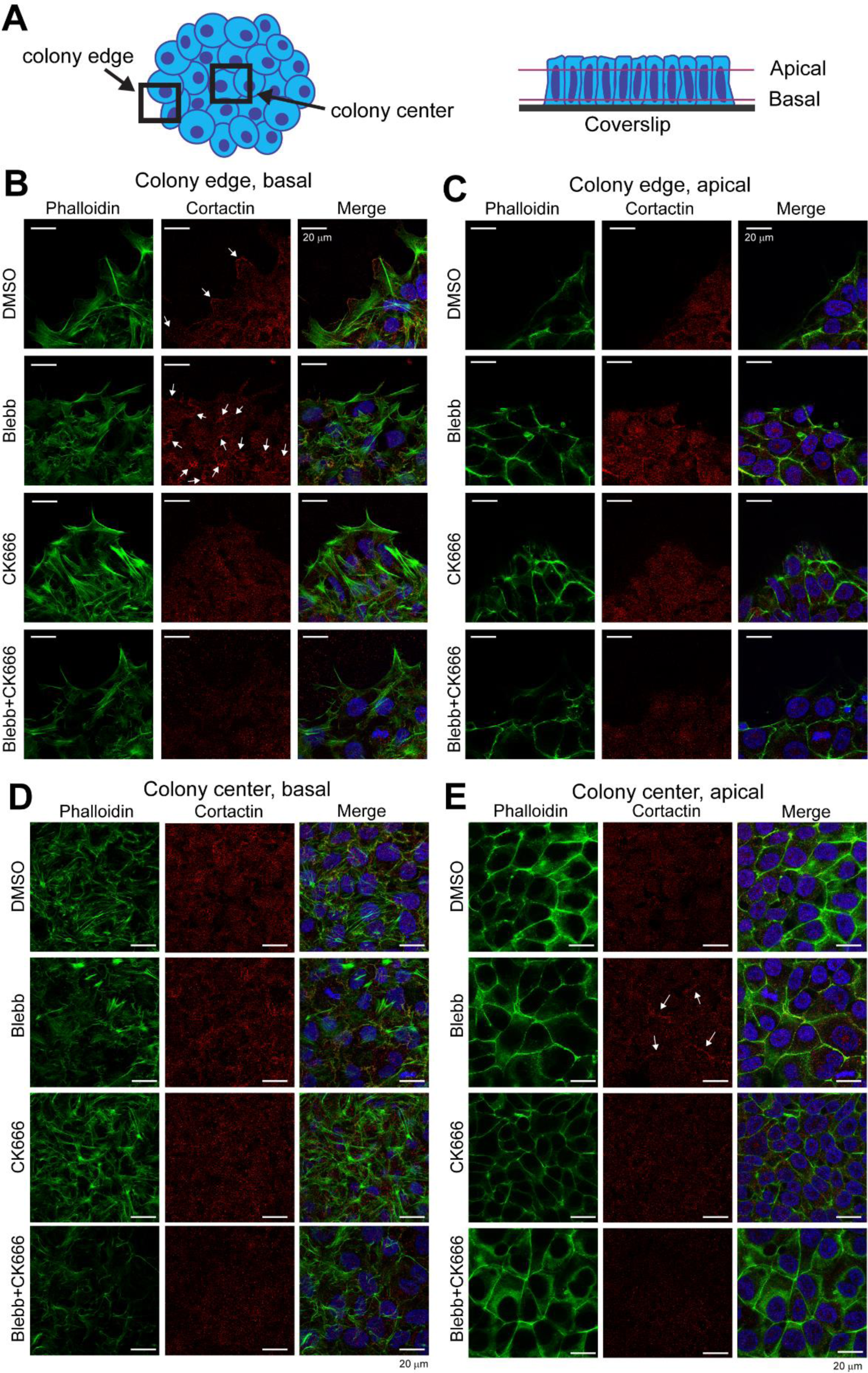
Stem cell colony edges possess cortactin decorated lamellipodia. **(A)** Illustration for stem cell colony imaging at the edge and centers. **(B-E)** Confocal z-stacks (63x/1.4 oil) were taken at H7-hESC colony edges and centers. Representative confocal immunofluorescence images of F-actin and cortactin after 3h DMSO, Blebbistatin (Blebb), CK666, or Blebb+CK666 treatments. **(B, D)** Shown are images at the bottom of the cells (basal) of the colony edge or center. **(C, E)** Images at the apical surface of colony at the edge or center. Arrows indicate cortactin decorated lamellipodia.

Since lamellipodia is a dynamic membrane structure regulated by Arp2/3 mediated dynamic actin meshwork, we asked if hPSCs at the colony edges possess dynamic lamellipodia with Arp2/3 at the leading edge. To test this, we co-transfected stem cell colonies with cationic liposomes containing GFP-ARP3 (Addgene #8462) (Welch et al., 1997) and CAG-mCherry (Addgene #108685) (Mishra et al., 2019), and performed live cell confocal imaging under control or inhibitory small molecules as explained in Figure 3A. Under the untreated condition, as well as in blebbistatin treated myosin II inhibition, we found edge cells localize GFP-ARP3 at the leading-edge with dynamic lamellipodia (Fig. 3 B, C; video 1) and (Fig. 3 E, F; video 2) respectively. Lamellipodia of the edge cells remained highly dynamic under myosin II inhibition as shown by the kymographs (Fig. 3 D, G) with a moderate increase in the membrane protrusion/retraction rates (Fig. 3 K). Furthermore, Arp2/3 inhibition by CK666 led to the loss of GFP-ARP3 localization at the periphery of leading-edge cells and corresponding membrane dynamics (Fig. 3 H, I; video 3). This observation was validated by the kymograph analysis of cell edges (Fig. 3 J) and corresponding membrane protrusion/retraction rate measurements (Fig. 3 K). Thus, our data shows stem cells at the colony edges but not at centers possess dynamic lamellipodia structures which are dependent on Arp2/3, but independent of actomyosin contractility.

**Figure 3:**
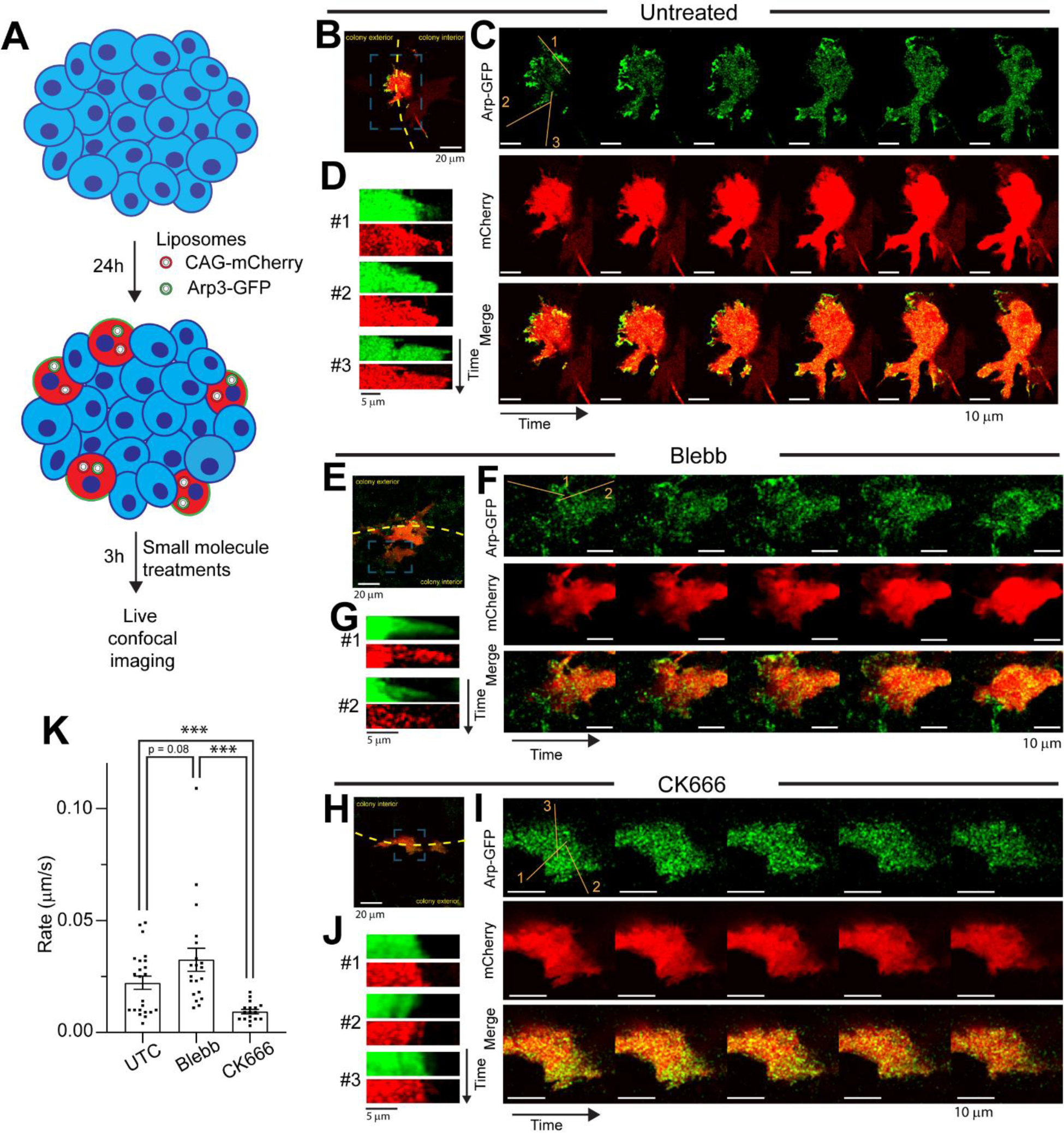
Lamellipodial dynamics of the edge cells is Arp2/3 dependent but independent of Myosin II activity. **(A)** Illustration of live cell imaging (63x/1.4 oil) of H7-hESCs 24h after dual transfection with cationic liposomes containing CAG-mCherry and GFP-ARP3 plasmids under indicated treatments. **(B, E, H)** Composite images show cell location relative to colony edge and center. **(C, F, I)** Montages of the timeseries taken with 15 second intervals for 5 minutes, showing edge dynamics of cells expressing ARP3 and mCherry from videos 1-3. **(D, G, J)** Kymographs corresponding to the numbered lines in the first frame of the montages. **(K)** Quantification of the movement rate (μm/sec) measured from the kymographs. Each data point represents a protrusion or retraction rate from the kymograph, n = 17-23 kymographs from 3-5 movies per treatment. Error bars are SEM. *** p < 0.001. Unpaired student’s *t-test* between independent datasets.

### Edge specific transfection is mitigated by loss of lamellipodia

Dynamic lamellipodia could provide negatively charged membrane surface for cationic liposome fusion and this process may be enhanced with more lamellipodia surface. To test this hypothesis, we first inhibited myosin II by blebbistatin which is known to increase lamellipodia membrane surface (Raucher and Sheetz, 2000) and then measured transfection efficiency by imaging and flow cytometry as explained in Figure 4 A. Flow gating strategy to count the percentage of the live singlet stem cell population expressing mCherry and single cell fluorescence intensity are described in Figure S3. We observed edge specific (blue arrows) as well as some inside the colony transfection (yellow arrows) under myosin II inhibition (Fig. 4 B). Cells inside the colony presumably got transfected through the lamellipodia like cortactin positive membrane that appeared at the edge and at the inside cells (Fig. 2 B) as well as at the apical surface of colony center under myosin II inhibition (Fig. 2 E). Remarkably, single cell analysis by flow showed a significant increase in the percentage of cells that got transfected with blebbistatin (Fig. 4 C, D). If more lamellipodia surface under myosin II inhibition allows cells to internalize more liposomes, this will lead to more CAG-mCherry plasmid delivery and increased mCherry expression. Indeed, we observed significant increase in fluorescence intensity per cell measured by flow (Fig. 4 E) or by confocal imaging (Fig. 4 F, G). Conversely, we tested if reducing lamellipodia by potent Arp2/3 inhibitor CK666 lowers liposome transfection efficiency. In agreement, we observed a strong reduction of cellular transfection efficiency measured by imaging and flow (Fig. 4 B, C, D). Furthermore, Arp2/3 inhibition reduced the amount of liposome delivery into cells as reveled by reduced fluorescence intensity per cell by flow (Fig. 4 E) and by confocal imaging (Fig. 4 F, G). The Arp2/3 inhibitory effect was to some extent rescued by myosin II inhibition, as increased transfected cell population (Fig. 4B, C, D) and cellular fluorescence intensity by flow (Fig. 4 E) and by imaging (Fig. 4 F, G) were observed when stem cell colonies were treated both by blebbistatin and CK666 compared to CK666 alone. Hence, this data suggests actomyosin contractility as an inhibitory factor, with Arp2/3 mediated dynamic lamellipodia facilitating the cationic liposome delivery into the cells.

**Figure 4:**
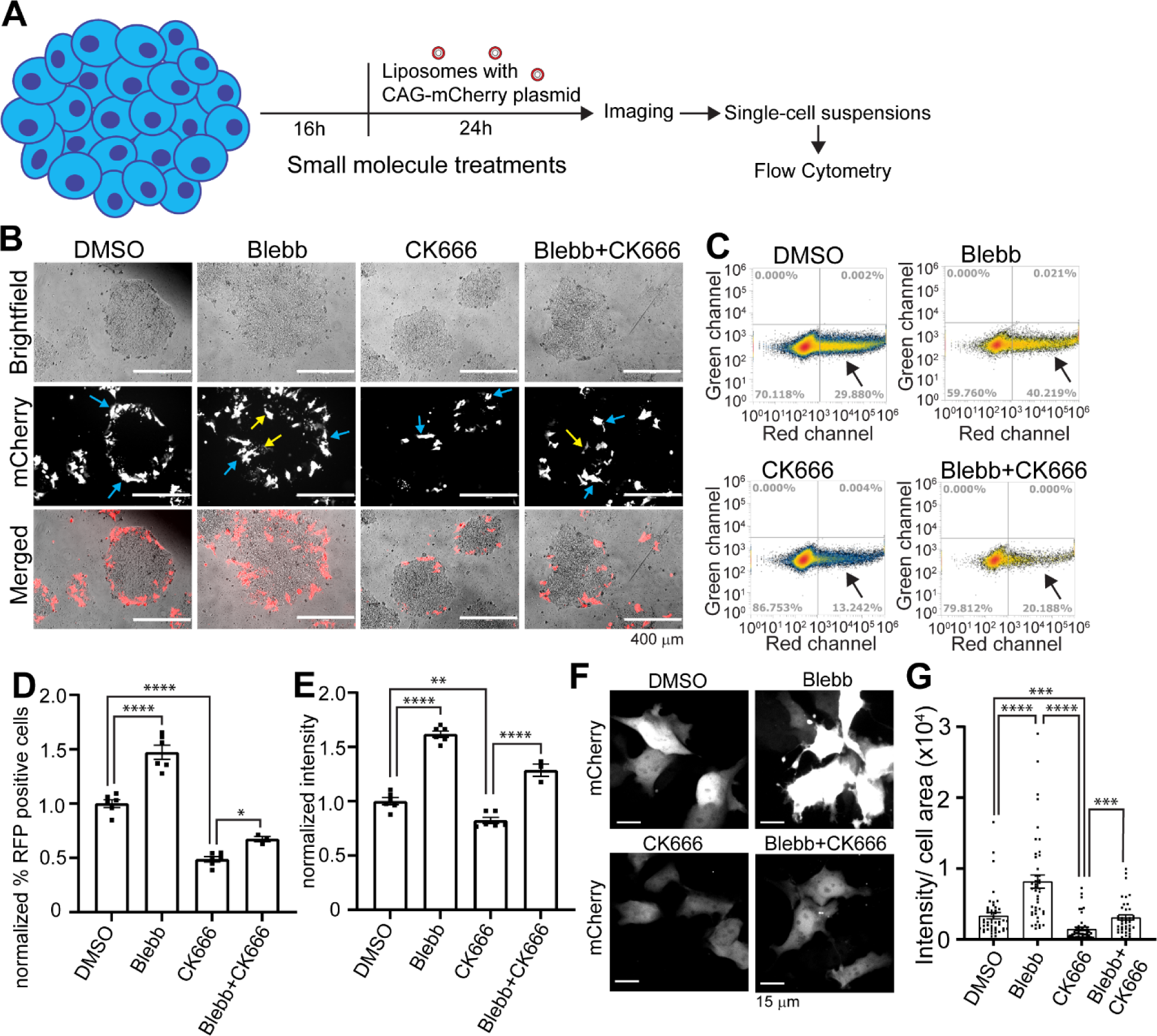
Edge specific transfection is dependent on Arp2/3 and augmented by Myosin II inhibition. **(A)** Illustration for stem cell colony treatment with small molecules and then transfection with cationic liposomes containing CAG-mCherry plasmid to measure transfection efficiency. **(B)** Representative images of stem cell colonies 24h post transfection in presence of small molecules. Blue arrows indicate transfected edge cells, yellow arrows indicate transfected cells inside the edges. **(C)** Single cell suspensions were collected 24h after transfection with indicated treatments and run through flow cytometer. RFP positive cell populations indicated by arrows quantified for **(D)** percentage of RFP-positive cells and **(E)** average mCherry intensity, normalized to DMSO. Each point represents an independent biological repeat. n=3-6. **(F)** Representative confocal images of H7-hESCs treated for 16h and then transfected with CAG-mCherry for 24h with continued treatment (63x/1.4 oil). **(G)** Quantification of mCherry intensity per cell area from sum projections of confocal z-stacks. Each point represents one cell, n = 40-47 cells. Error bars are SEM. * p < 0.05, ** p < 0.01, *** p < 0.001, **** p < 0.0001. Unpaired student’s *t-test* between independent datasets.

Given that the loss of myosin II contractility enhanced edge cell transfection, we next asked whether the process is actin cytoskeleton dependent. To test this, we first checked the minimum liposome exposure time needed to have successful transfection and how long cells can remain attached under actin cytoskeleton disruption. Based on a time course experiment of liposome transfection, we found that 1h exposure is sufficient for stem cell transfection. Colonies exposed to cationic liposomes containing mCherry plasmid for 1h to 24h as explained in Figure S4 A showed similar edge specific transfection (Fig. S4 B) with similar levels of percentage of cell transfection as measured by flow (Fig. S4 C-D). Cellular fluorescence intensity showed similar levels for all the different transection durations except for 24h which showed a moderate increase, suggesting that the number of liposomes entering into an individual cell is independent of liposome exposure time (Fig. S4 E). Furthermore, we found when actin cytoskeleton is disrupted by a potent actin depolymerizing drug Latrunculin A (Lat A) (Fujiwara et al., 2018) for 24h (Fig. S5 A), it led to a reduced number of live cells (Fig. S5 B, C, yellow gate), with a corresponding reduced number of mCherry expressing cells (Fig. S5 D, arrow). Prolonged Lat A treatment led to stem cell detachment from the dish as shown by reduced total cell count by flow (Fig. S5 E). So, we had to disrupt actin filaments for a small period of time followed by transfection to assess the role of actin in cationic liposome transfection. Based on these findings we disrupted actin cytoskeleton by Lat A for 1h and then exposed stem cell colonies for one more hour with liposomes containing CAG-mCherry plasmid along with Lat A, followed by exchanging to media without Lat A and liposomes, and measuring transfection after 24h in stem cell media as explained in Fig. 5 A. We saw actin cytoskeleton disruption did not inhibit cell transfection at the edge cells (Fig. 5 B). Interestingly, we saw a moderate increase in the percentage of transfected population measured by flow (Fig. 5 C, D) but with mild reduction in cellular fluorescence intensity (Fig. 5 E). These data support the prior findings where disruption of actin filaments showed increased direct penetration of cationic liposomes through plasma membrane fusion (Zhou and Huang, 1994). Reduction of cellular fluorescence intensity for Lat A treated colonies could be due to uptake of a smaller number of liposomes, limited by reduced lamellipodia surface at the edge cells. Lat A treatment for 2h followed by a wash did not detach cells as shown by imaging (Fig. 5 B) and by total cell count by flow after 24 (Fig. 5 F). Lat A treatment was effective in disrupting actin cytoskeleton within 3h as shown by confocal imaging of phalloidin labelled F-actin filaments (Fig. 5G). These data support the notion that cationic liposome transfection to cells depends on the availability of dynamic lamellipodial plasma membrane independent of actin cytoskeleton and myosin II activity.

**Figure 5:**
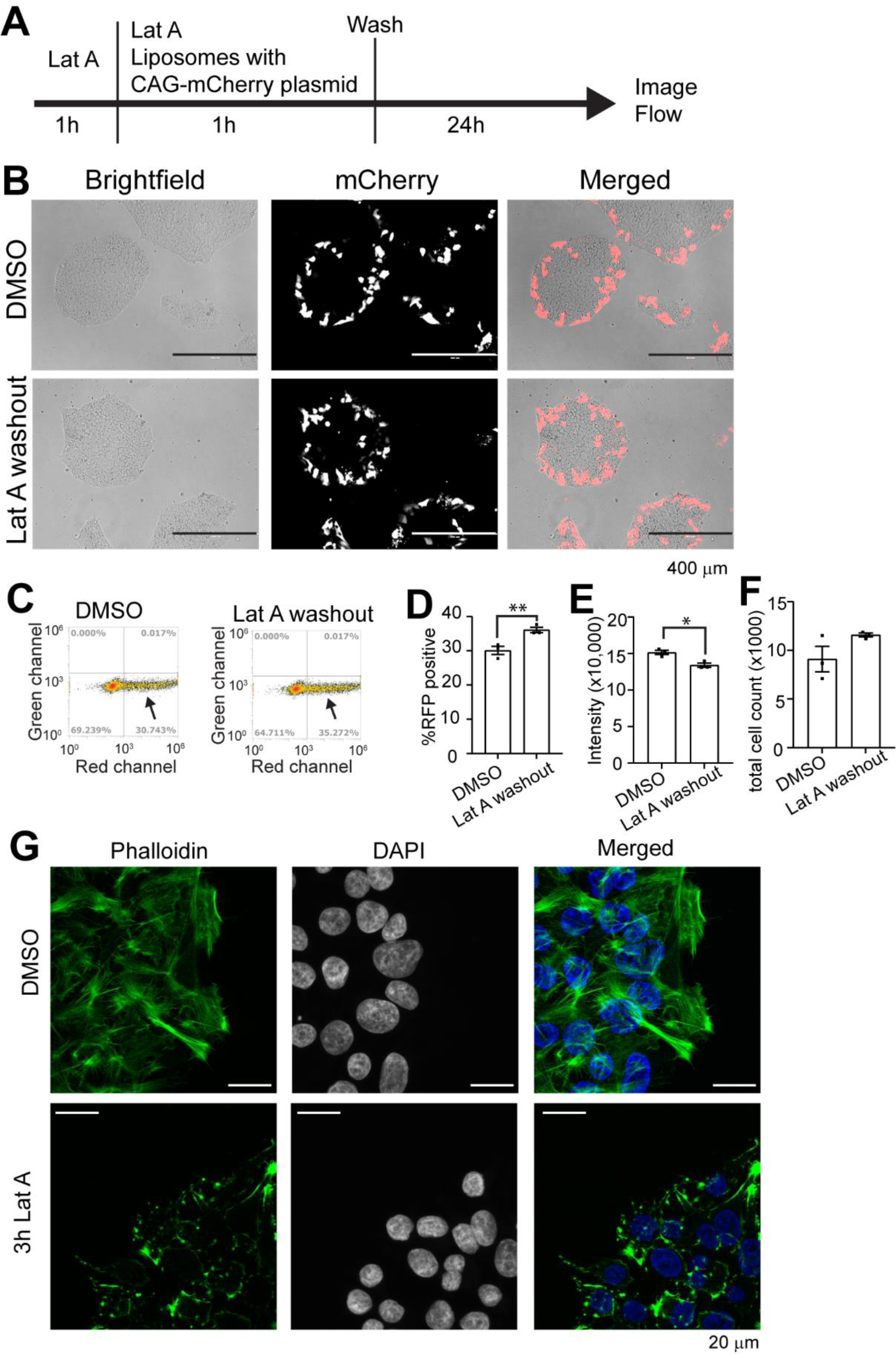
Edge transfection by cationic liposomes is independent of actin cytoskeleton integrity. **(A)** Illustration of stem cell colony transfection under Lat A treatments. H7-hESC colonies were pretreated with Lat A (2 μM) for 1h and then transfected for 1h with cationic liposomes containing CAG-mCherry plasmid along with Lat A. Media was then changed to remove plasmids and Lat A. **(B)** Images show edge specific transfection 24h after the Lat A washout. (**C**) Flow cytometry analysis on RFP positive cells (arrow) 24h after Lat A washout for **(D)** percentage of RFP positive cells, **(E)** mean RFP intensity per cell, and **(F)** total cell count. **(G)** Colonies were treated with Lat A for 3h, then fixed and stained for phalloidin to visualize the actin changes (63x/1.4 oil). Each point represents an independent biological repeat, n=3. Error bars are SEM. * p < 0.05, ** p < 0.01. Unpaired student’s *t-test* between independent datasets.

### Epithelial cell junctions limit cationic liposome transfection

While our data suggest that dynamic lamellipodia at the edge cells promote cationic liposome transfection, it is still not clear why the epithelial-like cells at the stem cell colony center are not transfected. It has been reported that the basement membrane of epithelial cells forms dynamic membrane structures with some similarity to lamellipodia (Nelson and Larsen, 2015). We asked if the cell junctions at the apical side limit liposome access to the basal membrane, and hence their transfection. So, we hypothesized that if we break these cell junctions that should promote transfection of the center cells. To test this, we enzymatically digested the cell surface proteins with either accutase or trypsin for a brief 1min or 3min and then transfected with cationic liposomes containing CAG-mCherry plasmid as explained in Fig. 6 A, B. Remarkably, we saw a significant increase in transfection inside the colony (Fig. 6 C). Short exposure of these enzymes maintained colony morphology with cells still attached to the coverslip (Fig. 6 C, D). But when we checked junctional integrity by staining against actin and tight junction protein Zonula occludens (ZO-1), we saw the opening of junctions inside the colony (Fig. 6 E, arrows). This further supports the notion that cell junction integrity limits cationic liposomes from accessing the basement membrane of the epithelial-like cells required for transfection. Since we observed some transfection inside the colony under myosin II inhibition by blebbistatin, we asked if this is due to the disrupted cell junctions. However, actin filament and ZO-1 staining revealed myosin II inhibition by blebbistatin does not compromise junctional integrity of the stem cell colony (Fig. 6 F). Thus, our data suggest cationic liposome delivery into the epithelial-like cells requires destabilization of the cellular junctions.

**Figure 6:**
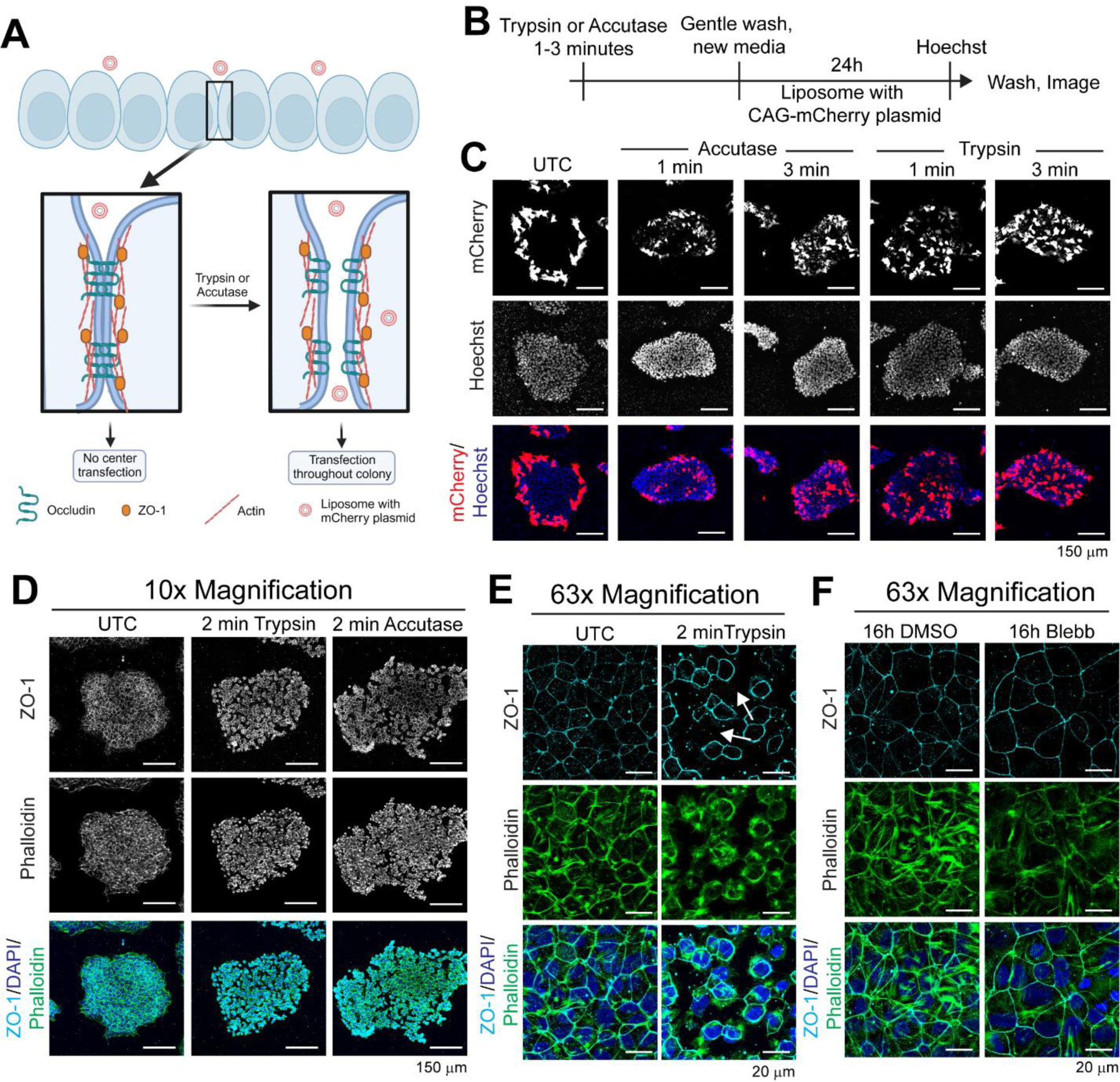
Junctional integrity of the epithelial like stem cell colony center inhibits cationic liposome transfection. **(A)** Experimental design to test if intact cell-cell junctions at the stem cell colony centers limit cationic liposome transfection. ZO-1 containing tight junctions are disrupted by trypsin/accutase. **(B)** H7-hESC colonies were exposed to trypsin or accutase for 1-3 minutes, washed, and then transfected with cationic liposomes containing CAG-mCherry plasmid. **(C)** Confocal images (10x/0.3) show untreated cells (UTC) with edge specific transfection, while colonies briefly treated with accutase, or trypsin got transfected both at the edge and center. **(D)** Images of stem cell colonies (10x/0.3) treated with trypsin or accutase for 2 min and then fixed and stained for ZO-1. **(E, F)** Max projections of confocal z-stacks (63x/1.4 oil) of ZO-1 in stem cell colony centers after **(E)** 2 min trypsin or **(F)** 16h blebbistatin treatments.

### Liposome delivery to epithelial cells require receptor mediated endocytosis

Liposome mediated therapeutic delivery to the epithelial cells through disrupting epithelial integrity is not ideal, as that would be associated with many health complications. Hence, we asked if these cells could take up liposomes through receptor mediated endocytosis without the need for disrupting epithelial integrity. To this end, we have used liposomes (m-Fluoroliposome-DiO, Encapsula) containing 4-aminophenyl α−D-mannopyranoside conjugated lipids and lipophilic water insoluble DiO dye whose excitation/emission peaks fall at 484/501 nm. 4-aminophenyl α−D-mannopyranoside is a hydrophobic derivative of mannose and can be internalized through the mannose receptor for drug delivery as illustrated in Fig. 7A. We used these liposomes to transfect stem cell colonies under control or small molecule treatments for 24h and measured transfection efficiency by imaging and flow as explained in Fig. 7 B. Much to our surprise, we see liposome transfection throughout the colony (Fig. 7 C). The colony center shows less fluorescence signal than the edge cells (Fig. 7 C), indicating that edge cells uptake liposomes through both membrane fusion and receptor mediated endocytosis. When we checked cells at the edge and center as illustrated in Fig. 7 D using high-magnification (63x) confocal imaging, we see robust DiO fluorescence signals, notably on the internal membranes for these cells (Fig. 7 E, F). As receptor mediated endocytosis depends on acto-myosin contractility (Chandrasekar et al., 2014; Wayt et al., 2021), we checked if the transfection efficiency is reduced under myosin II and Arp-2/3 inhibition by flow analysis as illustrated in Fig. 7 G. Indeed, we observed significantly reduced transfected cell populations under blebbistatin treatment. However, the effect was less under Arp-2/3 inhibition by CK666, but in presence of both blebbistatin and CK666 we saw maximum reduction (Fig. 7 H, I). Although, we see reduction in transfected cell populations in the presence of CK666 and blebbistatin, cells were still able to uptake some liposomes as revealed by DiO signals on images (Fig. 7 E, F) and by flow analysis (Fig. 7 H, I). This suggests that myosin II contractility on Arp-2/3 nucleated actin filaments aids in receptor mediated endocytosis of liposomes on the apical surface of the epithelial-like stem cells, with the existence of alternative mechanisms. It has been reported that viral transduction to cells occurs through receptor mediated endocytosis (Cossart and Helenius, 2014). To test if viral transduction would occur throughout the stem cell colonies, we transfected the stem cell colonies with lentivirus containing GFP vector under EF1α-short (EFS) promoter. Indeed, similar to the mannosylated liposomes, we observed stem cells throughout the colony got transduced with virus as shown by GFP expressions (Fig. S6).

**Figure 7:**
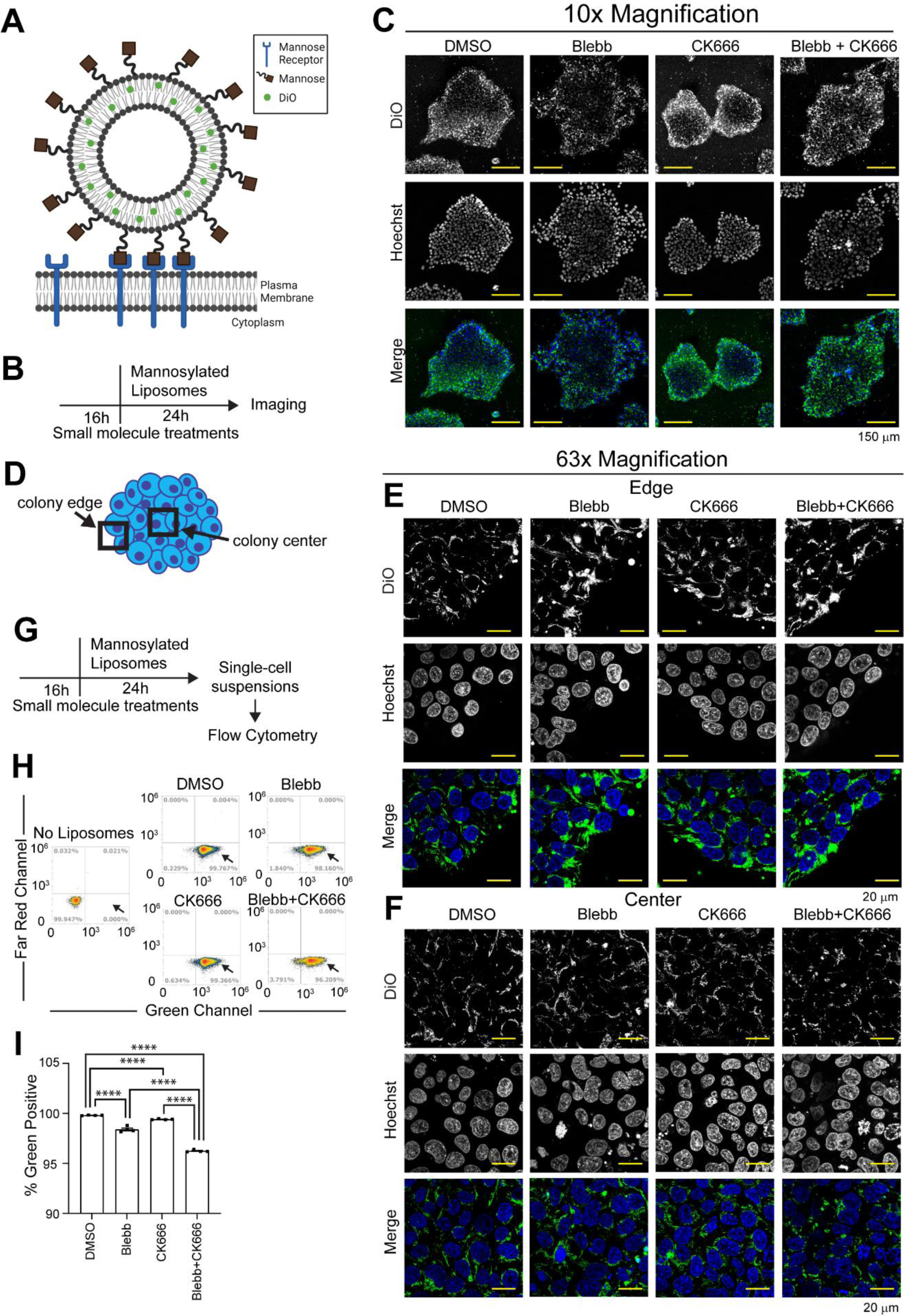
Epithelial like stem cell colony center internalizes liposomes through receptor mediated endocytosis. **(A)** Potential mechanism for receptor mediated liposome endocytosis where liposomes contain mannosylated lipids and DiO lipophilic dye. **(B)** Colonies were pre-treated with small molecules for 16h and then liposomes were added for another 24h before imaging. **(C)** Representative low magnification (10x/0.3) images of drug treated stem cell colonies 24h after adding mannosylasted liposomes. **(D)** High magnification (63x/1.4 oil) images were taken at the colony **(E)** edge and **(F)** center showing internalization of mannosylated liposomes. **(G)** Colonies were collected as single-cell suspensions and run through flow cytometry after indicated treatments. **(H)** Flow plots and quantification of the **(I)** percentage of green positive cells. Each point represents an independent biological repeat, n=3. Error bars are SEM. **** p < 0.0001. Unpaired student’s *t-test* between independent datasets.

Thus, our study here reveals unique mechanisms where Arp-2/3 mediated dynamic lamellipodia could be enhanced for increasing cationic liposome delivery to the migratory cells such as immune and cancer cells. However, receptor mediated endocytosis could be leveraged to promote liposome delivery to the epithelial cells. As a majority of the cell types in our body are either epithelial or migratory in nature, this study provides a foundation for increasing liposome based therapeutic delivery efficacy which is a safer treatment vehicle than virus.

## Discussion

We have shown here that human pluripotent stem cell colonies possess unique morphological features with cells at the colony edges showing dynamic lamellipodia but with cells at the center forming epithelial junctions with characteristic honeycomb shaped actin filament and junctional protein ZO-1 localizations at the apical side. We found cationic liposomes, which are a preferred vehicle for DNA/RNA delivery, preferentially delivered plasmids to the stem cell colony edge cells but not to the center. We further discovered cell junctions at the colony centers block such transfection, but mild enzymatic digestion of the cell surface proteins led to the transfection of entire colony. However, liposomes with mannosylated lipids are capable of transfecting the entire colony through receptor mediated endocytosis without the need for disrupting cell-cell junctions. These results are very important as they provide dual mechanisms for delivering liposomes to migratory as well as to epithelial cells as illustrated in our graphical abstract.

Our discovery that myosin II inhibition resulted in more cortactin decorated lamellipodia in the edge cells with enhanced liposome delivery to individual cell as well as to a greater number of cells within the colony is supported by existing literature. It has been reported that acto-myosin contractility negatively regulates lamellipodia formation (Raucher and Sheetz, 2000) and inhibition of myosin II contractility increases liposome transfection to stem cells (Yen et al., 2014). Hence, under myosin II inhibition more lamellipodia surface at the edge cells are likely to promote more liposome uptake through membrane fusion. Cationic liposomes could be internalized either through membrane fusion (Cypionka et al., 2009) or through endocytosis (Zabner et al., 1995; Zelphati and Szoka, 1996). It is unlikely that the colony edge specific transfection that we observed is through endocytosis which is largely dependent on myosin II activity (Chandrasekar et al., 2014; Wayt et al., 2021), but we saw enhanced edge transfection under myosin II inhibition. However, at this point we cannot rule out a myosin II independent but other molecular motor dependent endocytic uptake mechanism in the edge cells.

Our observation that the cationic liposomes are restricted from transfecting the epithelial like stem cell center cells is intriguing. This further supports a non-endocytic delivery mechanism for cationic liposomes, as otherwise they would have transfected the entire colony. On the contrary, dissociation of cell-cell junctions by mild enzymatic digestions led to robust center cell transfection by cationic liposomes, supporting the notion that liposomes need access to the dynamic basement membrane for membrane fusion and internalization. However, dynamic cells at the colony edges and epithelial-like cells at the centers are competent for endocytic uptake as we observed robust transfection of both cell types by the mannosylated liposomes, which is dependent on the mannose receptor mediated endocytosis. Mannosylated liposomes have also been shown as a robust delivery vehicle for mouse brain tissue (Umezawa and Eto, 1988).

Thus, our discovery here provides many fold applications for developing novel therapy delivery and disease modeling research. The identification of receptor mediated mannosylated liposome delivery throughout the human stem cell colony is very important. Gene editing such as by CRISPR/Cas9 in hPSCs provides a unique opportunity for disease modeling research. Gene-edited stem cells can be differentiated to the target cell types for understanding disease mechanisms for developing therapy. Conversely, patient derived induced pluripotent stem cells (iPSCs) with gene mutations can be corrected by CRISPR/Cas9 followed by differentiation to the effected cell type and injection for regenerative medicine (Mollashahi et al., 2023). While these strategies have tremendous clinical benefit and are currently under active research, gene-editing in hPSCs is an extremely inefficient process owing to their poor transfection property (Yang et al., 2013). Our discovery that accessing mannose receptor mediated endocytosis through mannosylated liposomes provided nearly 100% hPSC transfection throughout the colony can significantly increase stem cell editing rate by increasing the delivery efficiency of CRISPR/Cas9 nucleotides. In addition to direct application for stem cell research, our findings that Arp2/3 mediated lamellipodia promotes cationic liposome delivery to the migratory cells is very important. This gives an opportunity to specifically enhance DNA or RNA delivery to migratory cells such as immune cells for vaccination or to cancer cells for inducing apoptosis and selective elimination. Our discovery that the receptor-mediated endocytosis provides high-efficiency liposome delivery to epithelial cells is also clinically important as a majority of our tissues are epithelial in nature and could be targeted through these mechanisms.

## ACKNOWLEDGEMENTS

This work was supported by grant from the NIH, United States (R00EY028223) and startup package to Dr. Das from the Indiana University School of Medicine. We thank Dr. Donald Zack for kindly providing the H7-hESC, and EP1-iPSC human pluripotent stem cell lines. We thank Dr. Padmanabhan Pattabiraman for sharing the ZO-1 antibody.

## AUTHOR CONTRIBUTIONS

M.S. and A.D. designed the experiments, analyzed data and wrote manuscript; M.S. performed experiments and analyzed data with the help of K.A; A.D. conceived and supervised the project and revised the manuscript.

## Resource and Data Availability

### Lead Contact

Further information and requests for resources and reagents should be directed to Arupratan Das.

Stem cells and plasmids are available from the lead contact’s laboratory upon request and completion of the Material Transfer Agreement. Data are available in the article itself and its supplementary materials.

## EXPERIMENTAL MODEL AND SUBJECT DETAILS

### Quantification and Statistical analysis

All data presented are mean ± SEM. Cells were treated with small molecules at different time points as independent biological samples in independent culture wells. Statistical tests between two independent datasets were done by unpaired student’s *t-test*. Graphs were made using GraphPad Prism 9.0 software. Figures were made in Adobe Illustrator and cartoon illustrations were made using BioRender.com.

## METHOD DETAILS

Reagent list with catalog numbers is available in the resource table (Table1).

### Stem cell culture

H7-ESCs (WiCell, https://www.wicell.org/) and EP1-iPSCs (Bhise et al., 2013) were grown in mTeSR1 media (mT) in 5%CO_2_, 37°C incubator on matrigel (MG) coated plates. To obtain hPSC colonies, cells were passaged by clump passaging using Gentle Cell Dissociation Reagent (GD) after reaching 80% confluency. GD was added to cells for 4 min at 37°C, aspirated, then mT was used to resuspend colonies; cell suspensions were mixed by pipetting 3-4 times to break up the colonies into small clumps and then seeded into new MG coated wells. Clump passaged colonies were cultured for an additional 2-3 days before experiments. For single cell passaging, cells were incubated with accutase for 10 min and then quenched with double volume of mT with 5 μM blebbistatin (blebb). These cells were pelleted by centrifugation at 150G for 5 min, and resuspended in media with blebb, counted, and seeded at a density of 25,000/well of a 24-well plate.

### Plasmid amplification and purification

100 ng of the plasmid (CAG-mCherry or GFP-ARP3) was added to 50 μl of Top10 *E coli* and kept in ice for 5 min. CAG-mCherry was a gift from Jordan Green (Addgene plasmid # 108685; http://n2t.net/addgene:108685; RRID:Addgene_108685) and pEGFP-N1-ARP3 from Matthew Welch (Addgene plasmid # 8462; http://n2t.net/addgene:8462; RRID:Addgene_8462). The bacteria were then heat shocked to promote uptake of the plasmid at 42°C for 45 seconds before being placed back into ice for 2 min. 250 μl SOC media was added to the bacteria and incubated in a 37°C shaker for 1h before being added to 5 ml LB-broth with Carbenicillin (50 μg/ml, for CAG-mCherry) or Kanamycin (50 μg/ml, for GFP-ARP3) and incubated overnight at 37°C in a shaking incubator. Plasmid was extracted following the kit (Zymo D4210) protocol, and concentration was measured using nanodrop.

### Stem cell transfection

Human pluripotent stem cells (hPSCs) were cultured as described above. Single cells after accutase passage were transfected 24h after seeding. The clump passaged colonies were added to a larger volume of media and equally split into the wells of 24-well plates. Cells were cultured for another 2-3 days until the colonies were established with distinct edges and centers, having a colony size around 1/10^th^ the size of a 10x objective field of view size at start of drug treatment or transfection. 24h after transfection, images were taken by the EVOS fluorescence microscope (Thermo Fisher Scientific) and cells were collected for flow cytometry. Using ImageJ software, fluorescence intensity was quantified by drawing a ‘donut’ containing the colony edge, measured as the edge; the ‘hole’ of the donut was then measured as the center. Raw integrated density was divided by the total area to get the average intensity per area for both edge and center of each colony.

Colonies were treated with 5 μM blebb, 100 μM CK666, both blebb and CK666, or the equivalent volume of DMSO in mT for the indicated time points and then transfected. Cell transfections (for 1 well of a 24well plate) were done by mixing 2 μl of lipofectamine stem (Invitrogen) and 600 ng of indicated plasmids in 50 μl optimem, vortexing, and letting the mixture sit at room temperature. 10 min after vortexing, this mixture was added by dropping onto the cell culture media and incubating for 24h.

Stem cell colonies were treated with Latrunculin A (Lat A) at 2 μM dose for a shorter timepoint of 1h and then transfected for another 1h as longer exposure (24h) detaches colonies. After which, the media with drug and liposomes was removed and the colonies incubated with fresh media for 24h.

### Flow Cytometry

Stem cell colonies after transfection for 24h were incubated in 30 μl accutase for 10 min, then quenched with 170 μl mT with 5 μM blebb. This 200 μl cell suspension was transferred into a 96-well round-bottom plate and read on the Attune NxT Acoustic Focusing Flow Cytometer (Thermo) equipped with Attune Auto Sampler (Thermo). Flow gating to separate live and dead cells was done by treating hPSCs with 2 μM puromycin, which induces cell death, or an equivalent amount of DMSO control for 48h and then made into a single cell suspension as described above. The cell population that disappeared under puromycin treatment was considered as the live cell population, while the remaining cells labelled with dead cell marker propidium iodide were considered as dead cells (Fig. S3 A, B). These gates were then used for the subsequent transfection experiments. From the live cell population, the singlet population was then gated, and from the singlet population, the percentage of RFP or green positive cells or the average single cell fluorescence intensities were measured using the Attune NxT Software. These gating conditions from each experiment are shown in supplemental figure 3. Data were exported to Excel or Prism for analysis and plotting. Each well was considered an independent biological sample, and three or more biological repeats were used for each condition.

### Immunofluorescence imaging

H7-hESCs were seeded using GD passaging on MG coated glass coverslips (1.5 thickness) and cultured for 2-3 days until colonies were established. Following the indicated treatments media was aspirated and cells were washed with 1X PBS, and then fixed with 4% Paraformaldehyde for 30 min at 37°C. At this step, if needed cells were washed once and then stored in PBS at 4°C until immunostained. Fixed cells were permeabilized with 0.5% Triton-X100 in PBS for 5 min and then washed in washing buffer (1% donkey serum, 0.05% Triton-X100 in PBS) 3 times for 5 minutes each. Cells were then incubated in blocking buffer (5% donkey serum, 0.2% Triton-X100 in PBS) for 1 hour at room temperature. After blocking, antibody against cortactin (Rabbit, Cell Signaling) or ZO-1 (Mouse, Invitrogen) was added (1:200 in blocking buffer) and the coverslips were incubated overnight at 4°C. Next, coverslips were washed with washing buffer 3 times for 5 minutes each and incubated for 2 hours at room temperature in the dark with secondary antibody anti-rabbit Alexa-568 (Invitrogen) or anti-mouse Alexa-647 (Invitrogen) (1:500 in blocking buffer), and Alexa Fluor 488 conjugated Phalloidin (4U/ml) for F-actin. The coverslips were washed with washing buffer 3 times for 5 minutes each, with 1.43 μM DAPI added to the second wash. Coverslips were then mounted using DAKO and sealed with clear nail polish. Confocal z-stack images were taken using Zeiss LSM700 with 63x/1.4 oil and 10x/0.3 air objectives.

### Live cell imaging

H7-hESCs were seeded into glass bottom MatTek dishes using GD passaging. After transfection, dishes were imaged using Tokai Hit live cell imaging chamber at 37°C and 5%CO_2_. Confocal z-stacks were taken using Zeiss LSM700 with 63x/1.4 oil objective and 10x/0.3 air objective. For lamellipodial measurements, colonies were transfected with cationic liposomes containing CAG-mCherry and GFP-ARP3 plasmids, using 600 ng of each plasmid. 24h post transfection, media with small molecules was added and incubated for 3h. Dishes were then placed into the Tokai Hit live cell chamber and a timeseries with 15 sec intervals for 5 min with simultaneous acquisition of red and green fluorescence signals. Lamellipodial dynamics were measured from kymographs using ImageJ software. A line was placed over the moving edge of GFP-ARP3 expressing cell and a kymograph made using the Kymograph plugin. Edge protrusion and retraction rates were quantified from kymographs for edge movement rates.

For single cell fluorescence intensity measurements, colonies were first treated with the indicated small molecules at the indicated timepoints, then transfected with cationic liposomes containing CAG-mCherry plasmid and incubated for another 24h in the presence of the small molecules before imaging. Confocal z-stacks of transfected cells were taken at the colony edges and intensity per cell area from the sum projections were measured using ImageJ software.

For partial digestion of the cell junctions (Fig 6), colonies were incubated with 200 μl accutase or trypsin for 1-3 minutes. Then trypsin or accutase was aspirated and cells gently washed twice with normal mT media. Treated colonies were then transfected with cationic liposome containing CAG-mCherry plasmid and incubated for 24h. Colonies were washed and incubated in mT with live cell nuclear dye Hoechst (5 μg/ml) for 15 min, washed, and then imaged using both 63x/1.4 oil and 10x/0.3 objectives.

### Stem cell transfection with mannosylated liposomes

Stem cell colonies in MatTek dishes were treated with the indicated small molecules or DMSO vehicle control at indicated time points, then 10 μL of liposomes (Encapsula Nanosciences) was added to the cell culture wells in the presence of small molecules and gently mixed by pipetting. After 24h incubation, media was aspirated, colonies were washed twice with mT, and then 2 mL of mT with 1 μL Hoescht (5 μg/ml) was added for live confocal imaging using both 63x/1.4 oil and 10x/0.3 objectives.

### Lentivirus

Stem cells at ∼80% confluency was clump passaged using GD and seeded into 96-well MG coated wells. The next day, cell counting was done from one well using accutase mediated single cell dissociation. Lentivirus (LV) (Life Technologies Cat # A32060) with viral vector containing P_EFS_-GFP was added to each well at a multiplicity of infection (MOI) of 10 along with 8 μg/mL polybrene. The plate was then centrifuged at 800 G at room temperature for 1h before placing it into a 37°C, 5% CO_2_ incubator overnight. The next day, media with LV was replaced with normal mT and GFP signal was observed over time with daily media exchange.

**Video 1:**

Confocal live cell imaging of untreated H7-hESCs 24h after dual transfection with CAG-mCherry and GFP-ARP3 plasmids (63x/1.4 oil). Movies acquired by taking images at 15 second intervals for 5 minutes.

**Video 2:**

Confocal live cell imaging of H7-hESCs after dual transfection with CAG-mCherry and Arp-GFP plasmids for 24h, followed by 3h treatment with Blebbistatin (63x/1.4 oil). Movies acquired by taking images at 15 second intervals for 5 minutes.

**Video 3:**

Confocal live cell imaging of H7-hESCs after dual transfection with CAG-mCherry and Arp-GFP plasmids for 24h, followed by 3h treatment with CK666 (63x/1.4 oil). Movies acquired by taking images at 15 second intervals for 5 minutes.

**Supplemental Figure S1: Low magnification images show edge specific CAG-mCherry colony transfection.**

Edge specific transfection can be seen in multiple H7-hESC colonies at 4x magnification 24h after transfection with cationic liposomes containing CAG-mCherry plasmid.

**Supplemental Figure S2: Edge specific transfection of hPSC colonies is independent of stem cell type.**

EP1-iPSCs were transfected with cationic liposomes containing CAG-mCherry plasmid. **(A)** Representative brightfield and fluorescence images of EP1-iPSCs after 24h transfection. **(B)** Fluorescence intensity trace through the colony center and edge as shown in the blue line in A. **(C)** Quantification of fluorescence intensity of colony edges and centers. Error bars are SEM. Each data point represents a colony, n=13. Unpaired student’s *t-test*, *** p < 0.001.

**Supplemental Figure S3: Flow Cytometry gating strategy**

Treated and transfected H7-hESCs were made into single-cell suspensions and run through flow cytometry. To create the gates for the living and dead cell populations, H7-hESCs were treated with 2 μM puromycin for 48h to induce cell death and then stained with propidium iodide which labels dead cells. **(A)** The controls show populations of both living (yellow gate) and dead (red gate) cells while **(B)** puromycin only has a dead cell population. These gates were then used for the subsequent transfection experiments to analyze live cell populations. From the live cells, the singlet population was then gated, and that singlet population further analyzed for percentage of cells expressing mCherry in the red channel or DiO in the green channel, as well as the average fluorescence intensity in single cells. (**C-G**) Gates are shown for the analysis done on indicated main or supplemental figures.

**Supplemental Figure S4: An hour exposure to cationic liposomes is sufficient for stem cell colony transfection.**

**(A)** Schematic of cationic liposomes containing CAG-mCherry plasmid transfection with different exposure time followed by liposome washout, media exchange and imaging 24h from the point liposomes were added. **(B)** Images show edge specific transfection of H7-hESCs from 1h to 24h liposome exposure times. (**C**) Single-cell suspensions were analyzed by flow cytometry showing the RFP positive cell distribution and the quantification of **(D)** percentage of RFP positive cells and **(E)** mean RFP intensity per cell. Each point represents an independent biological repeat, n=3. Error bars are SEM. * p < 0.05, ** p < 0.01, *** p < 0.001. Unpaired student’s *t-test* between independent datasets.

**Supplemental Figure S5: Prolonged disruption of actin cytoskeleton is toxic to stem cells.**

**(A)** Colonies were pretreated with Lat A (2 μM) for 16h and then transfected with liposome containing CAG-mCherry plasmid in the presence of Lat A followed by imaging and flow analysis. **(B)** Representative images 24h after transfection. Single-cell suspensions were then analyzed by flow cytometry showing the **(C)** live and dead cell population and **(D)** RFP positive cell distribution (arrow), and the quantification of **(E)** total cell count. Each point represents an independent biological repeat, n=3. Error bars are SEM. ** p < 0.01. Unpaired student’s *t-test* between independent datasets.

**Supplemental Figure S6: Cells throughout the hPSC colony are transduced by lentivirus.**

H7-hESCs were transduced with lentivirus containing GFP under P_EFS_ promoter. Shown are representative images at the indicated timepoints from 3 independent repeats.

## RESOURCES TABLE

| REAGENT or RESOURCE | SOURCE | IDENTIFIER |
| --- | --- | --- |
| Antibodies |  |  |
| Cortactin (H222) | Cell Signaling Technology | Cat# 3503 |
| ZO-1 Monoclonal Antibody (ZO1-1A12) | Invitrogen | Cat# 33-9100 |
| Alexa Fluor 488 Phalloidin | Invitrogen | Cat# A12379 |
| Donkey anti-Rabbit Alexa Fluor 568 | Invitrogen | Cat# A10042 |
| Goat anti-Mouse Alexa Fluor 647 | Invitrogen | Cat# 31541 |
| Bacterial and Virus Strains |  |  |
| Invitrogen™ One Shot™ TOP10 Chemically Competent *E. coli* | Fisher Scientific | Cat# C404003 |
| LentiArray™ CRISPR Positive Control Lentivirus, human HPRT, with GFP (P<sub>U6</sub>-HPRT1-P<sub>EFS</sub>-GFP) | ThermoFisher (Life Technologies) | Cat# A32060 |
| Cell Lines |  |  |
| H7-hESCs | WiCell; Sluch et al., 2018 |  |
| EP1-iPSCs | Bhise et al., 2013 |  |
| Chemicals and Reagents |  |  |
| Matrigel | Corning | Cat# CB40230 |
| Gentle Cell Dissociation Reagent | Stem Cell Technology | Cat# 7174 |
| mTeSR1 | Stem Cell Technology | Cat# 85850 |
| Accutase | Sigma | Cat# A6964 |
| TrypLE™ Express Enzyme (1X), phenol red (Trypsin) | ThermoFisher (Life Technologies) | Cat# 12605010 |
| Blebbistatin | Sigma | Cat# B0560 |
| CK666 | Sigma | Cat# SML0006 |
| Latrunculin A | ThermoFisher (Life Technologies) | Cat# L12370 |
| DAPI | Molecular Probes | Cat# D1206 |
| Hoechst 33342, Trihydrochloride, Trihydrate - 10 mg/mL Solution in Water | Invitrogen | Cat# H3570 |
| DMSO | Sigma | Cat# 276855 |
| Lipofectamine Stem | Invitrogen | Cat# STEM00003 |
| Optimem | Gibco | Cat# 31985070 |
| Paraformaldehyde 16% solution, EM grade | Electron Microscopy Sciences | Cat# 15710 |
| Triton-X-100 | Sigma | Cat# T8787 |
| Donkey Serum | Sigma | Cat# D9663 |
| SOC media | Fisher Scientific | Cat# BP974010X5 |
| Carbenicillin disodium salt | Sigma | Cat# C1389 |
| Kanamycin sulfate from *Streptomyces kanamyceticus* | Sigma | Cat# K0879 |
| LB Broth (Miller) | Sigma | Cat# L3522 |
| Polybrene | Sigma | Cat# TR-1003-G |
| ZymoPURE™ Plasmid Miniprep Kit | Zymo | Cat# D4210 |
| m-Fluoroliposome®-DiO | Encapsula NanoSciences | Cat# CLD-8925 |
| Plasmids |  |  |
| pCAG-mCherry plasmid | Addgene | Cat# 108685 |
| pEGFP-N1-ACTR3 plasmid (GFP-ARP3) | Addgene | Cat# 8462 |

| Software and Algorithms |  |
| --- | --- |
| ImageJ | NIH |
| Attune NxT Software | ThermoFisher |
| Prism version 9 | GraphPad |
| Zen Microscope Software | Zeiss |
| Adobe Illustrator | Adobe |
| BioRender | BioRender.com |

## REFERENCES

Bennett, C.F., M.Y. Chiang, H. Chan, J.E. Shoemaker, and C.K. Mirabelli. 1992. Cationic lipids enhance cellular uptake and activity of phosphorothioate antisense oligonucleotides. Mol Pharmacol. 41:1023–1033.

Bhise, N.S., K.J. Wahlin, D.J. Zack, and J.J. Green. 2013. Evaluating the potential of poly(beta-amino ester) nanoparticles for reprogramming human fibroblasts to become induced pluripotent stem cells. Int J Nanomedicine. 8:4641–4658.

Chandrasekar, I., Z.M. Goeckeler, S.G. Turney, P. Wang, R.B. Wysolmerski, R.S. Adelstein, and P.C. Bridgman. 2014. Nonmuscle myosin II is a critical regulator of clathrin-mediated endocytosis. Traffic. 15:418–432.

Cossart, P., and A. Helenius. 2014. Endocytosis of viruses and bacteria. Cold Spring Harb Perspect Biol. 6.

Cypionka, A., A. Stein, J.M. Hernandez, H. Hippchen, R. Jahn, and P.J. Walla. 2009. Discrimination between docking and fusion of liposomes reconstituted with neuronal SNARE-proteins using FCS. Proc Natl Acad Sci U S A. 106:18575–18580.

Doherty, G.J., and H.T. McMahon. 2009. Mechanisms of endocytosis. Annu Rev Biochem. 78:857–902.

Felgner, P.L., and G.M. Ringold. 1989. Cationic liposome-mediated transfection. Nature. 337:387–388.

Fujiwara, I., M.E. Zweifel, N. Courtemanche, and T.D. Pollard. 2018. Latrunculin A Accelerates Actin Filament Depolymerization in Addition to Sequestering Actin Monomers. Current Biology. 28:3183-3192.e3182.

Goeckeler, Z.M., P.C. Bridgman, and R.B. Wysolmerski. 2008. Nonmuscle myosin II is responsible for maintaining endothelial cell basal tone and stress fiber integrity. American Journal of Physiology-Cell Physiology. 295:C994–C1006.

Kelly, C., C. Jefferies, and S.A. Cryan. 2011. Targeted liposomal drug delivery to monocytes and macrophages. J Drug Deliv. 2011:727241.

Kim, Y., H. Jang, K. Seo, J.H. Kim, B. Lee, H.M. Cho, H.J. Kim, E. Yang, H. Kim, J.A. Gim, Y. Park, J.R. Ryu, and W. Sun. 2022. Cell position within human pluripotent stem cell colonies determines apical specialization via an actin cytoskeleton-based mechanism. Stem Cell Reports. 17:68–81.

Kovács, M., J. Tóth, C. Hetényi, A. Málnási-Csizmadia, and J.R. Sellers. 2004. Mechanism of Blebbistatin Inhibition of Myosin II*. Journal of Biological Chemistry. 279:35557–35563.

Lehtimäki, J.I., E.K. Rajakylä, S. Tojkander, and P. Lappalainen. 2021. Generation of stress fibers through myosin-driven reorganization of the actin cortex. eLife. 10:e60710.

Mishra, B., D.R. Wilson, S.R. Sripathi, M.P. Suprenant, Y. Rui, K.J. Wahlin, C.A. Berlinicke, J.J. Green, and D.J. Zack. 2019. A combinatorial library of biodegradable polyesters enables non-viral gene delivery to post-mitotic human stem cell-derived polarized RPE monolayers. Regen Eng Transl Med. 6:273–285.

Mollashahi, B., H. Latifi-Navid, I. Owliaee, S. Shamdani, G. Uzan, S. Jamehdor, and S. Naserian. 2023. Research and Therapeutic Approaches in Stem Cell Genome Editing by CRISPR Toolkit. Molecules. 28:1982.

Nelson, D.A., and M. Larsen. 2015. Heterotypic control of basement membrane dynamics during branching morphogenesis. Developmental Biology. 401:103–109.

Raucher, D., and M.P. Sheetz. 2000. Cell Spreading and Lamellipodial Extension Rate Is Regulated by Membrane Tension. Journal of Cell Biology. 148:127–136.

Sercombe, L., T. Veerati, F. Moheimani, S.Y. Wu, A.K. Sood, and S. Hua. 2015. Advances and Challenges of Liposome Assisted Drug Delivery. Front Pharmacol. 6:286.

Suraneni, P., B. Rubinstein, J.R. Unruh, M. Durnin, D. Hanein, and R. Li. 2012. The Arp2/3 complex is required for lamellipodia extension and directional fibroblast cell migration. Journal of Cell Biology. 197:239–251.

Umezawa, F., and Y. Eto. 1988. Liposome targeting to mouse brain: Mannose as a recognition marker. Biochemical and Biophysical Research Communications. 153:1038–1044.

Wayt, J., A. Cartagena-Rivera, D. Dutta, J.G. Donaldson, and C.M. Waterman. 2021. Myosin II isoforms promote internalization of spatially distinct clathrin-independent endocytosis cargoes through modulation of cortical tension downstream of ROCK2. Mol Biol Cell. 32:226–236.

Weed, S.A., and J.T. Parsons. 2001. Cortactin: coupling membrane dynamics to cortical actin assembly. Oncogene. 20:6418–6434.

Welch, M.D., A.H. DePace, S. Verma, A. Iwamatsu, and T.J. Mitchison. 1997. The human Arp2/3 complex is composed of evolutionarily conserved subunits and is localized to cellular regions of dynamic actin filament assembly. J Cell Biol. 138:375–384.

Wu, C., S.B. Asokan, M.E. Berginski, E.M. Haynes, N.E. Sharpless, J.D. Griffith, S.M. Gomez, and J.E. Bear. 2012. Arp2/3 is critical for lamellipodia and response to extracellular matrix cues but is dispensable for chemotaxis. Cell. 148:973–987.

Yang, L., M. Guell, S. Byrne, J.L. Yang, A. De Los Angeles, P. Mali, J. Aach, C. Kim-Kiselak, A.W. Briggs, X. Rios, P.-Y. Huang, G. Daley, and G. Church. 2013. Optimization of scarless human stem cell genome editing. Nucleic Acids Research. 41:9049–9061.

Yen, J., L. Yin, and J. Cheng. 2014. Enhanced Non-Viral Gene Delivery to Human Embryonic Stem Cells via Small Molecule-Mediated Transient Alteration of Cell Structure. J Mater Chem B. 2:8098–8105.

Zabner, J., A.J. Fasbender, T. Moninger, K.A. Poellinger, and M.J. Welsh. 1995. Cellular and molecular barriers to gene transfer by a cationic lipid. J Biol Chem. 270:18997–19007.

Zelphati, O., and F.C. Szoka. 1996. Mechanism of oligonucleotide release from cationic liposomes. Proceedings of the National Academy of Sciences. 93:11493–11498.

Zhou, X., and L. Huang. 1994. DNA transfection mediated by cationic liposomes containing lipopolylysine: characterization and mechanism of action. Biochim Biophys Acta. 1189:195–203.

