## Supplemental Figures for "Arp2/3 mediated dynamic lamellipodia of the hPSC colony edges promote liposome-based DNA delivery"

### Supplemental Figure S1

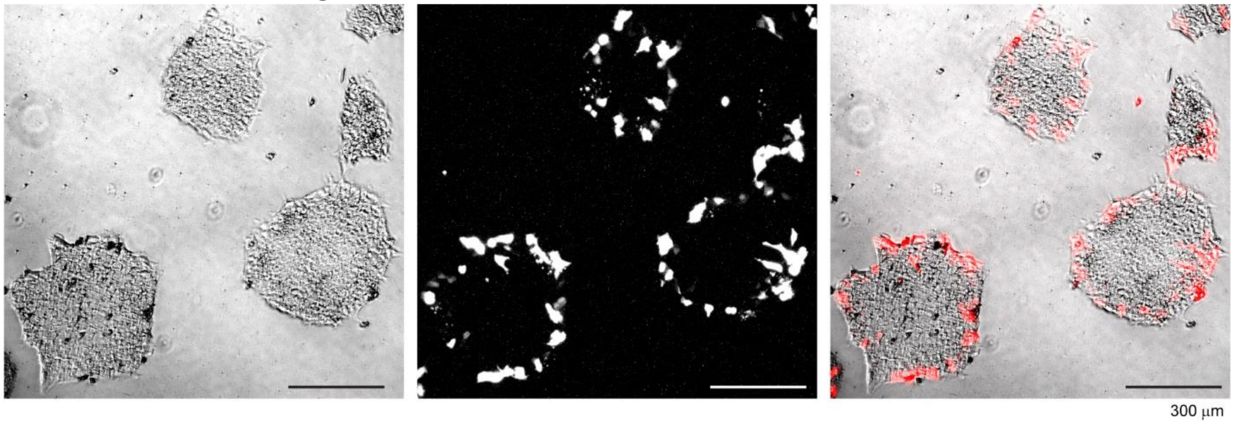

### Supplemental Figure S2

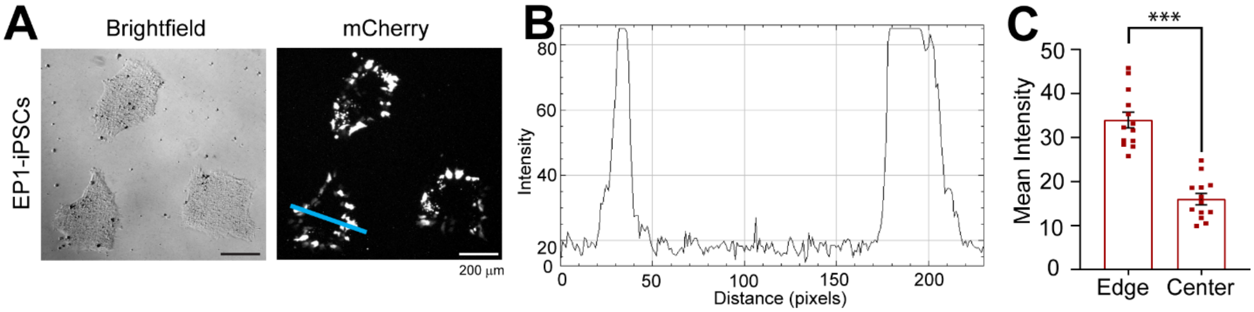

### Supplemental Figure S3

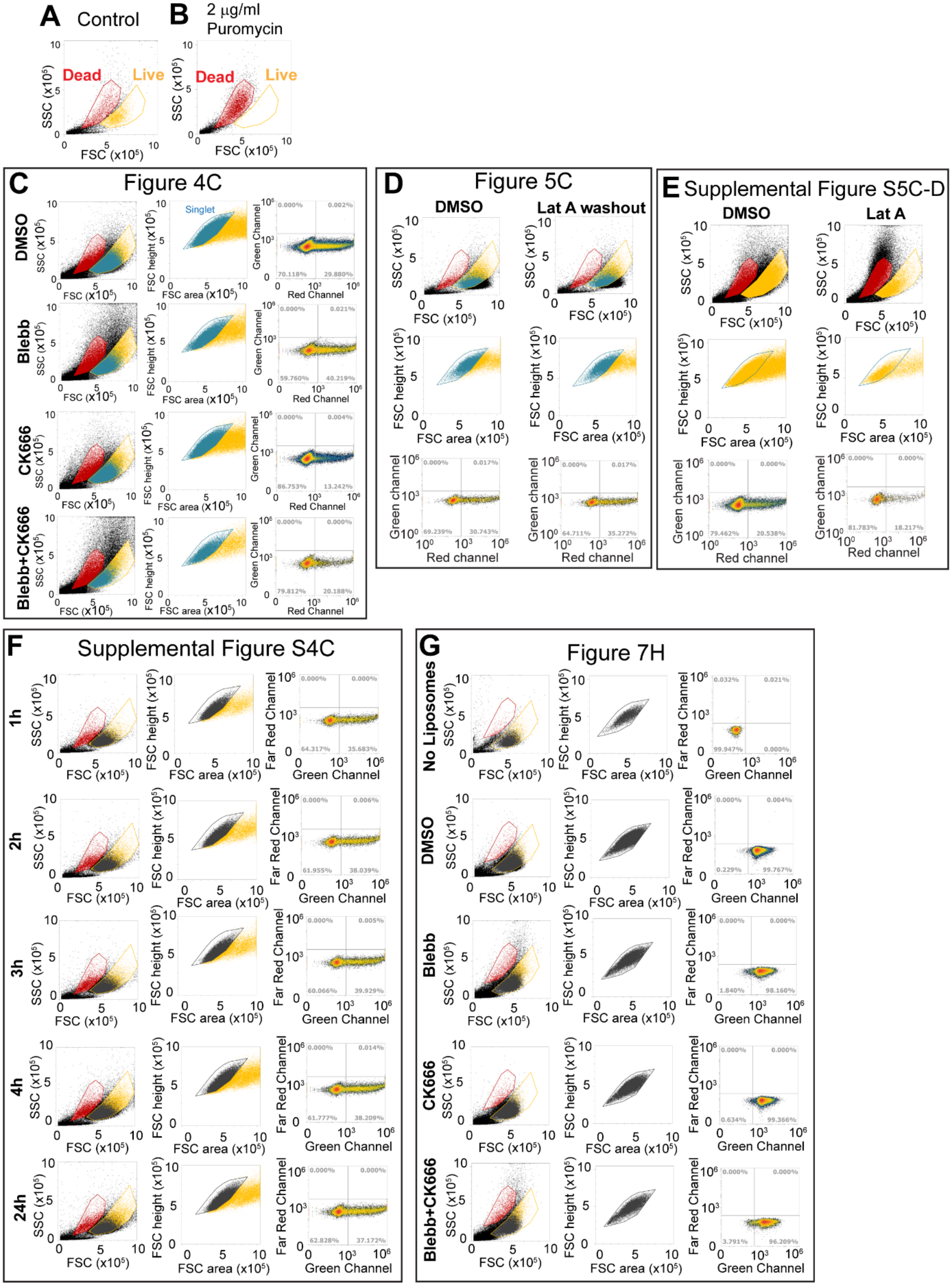

### Supplemental Figure S4

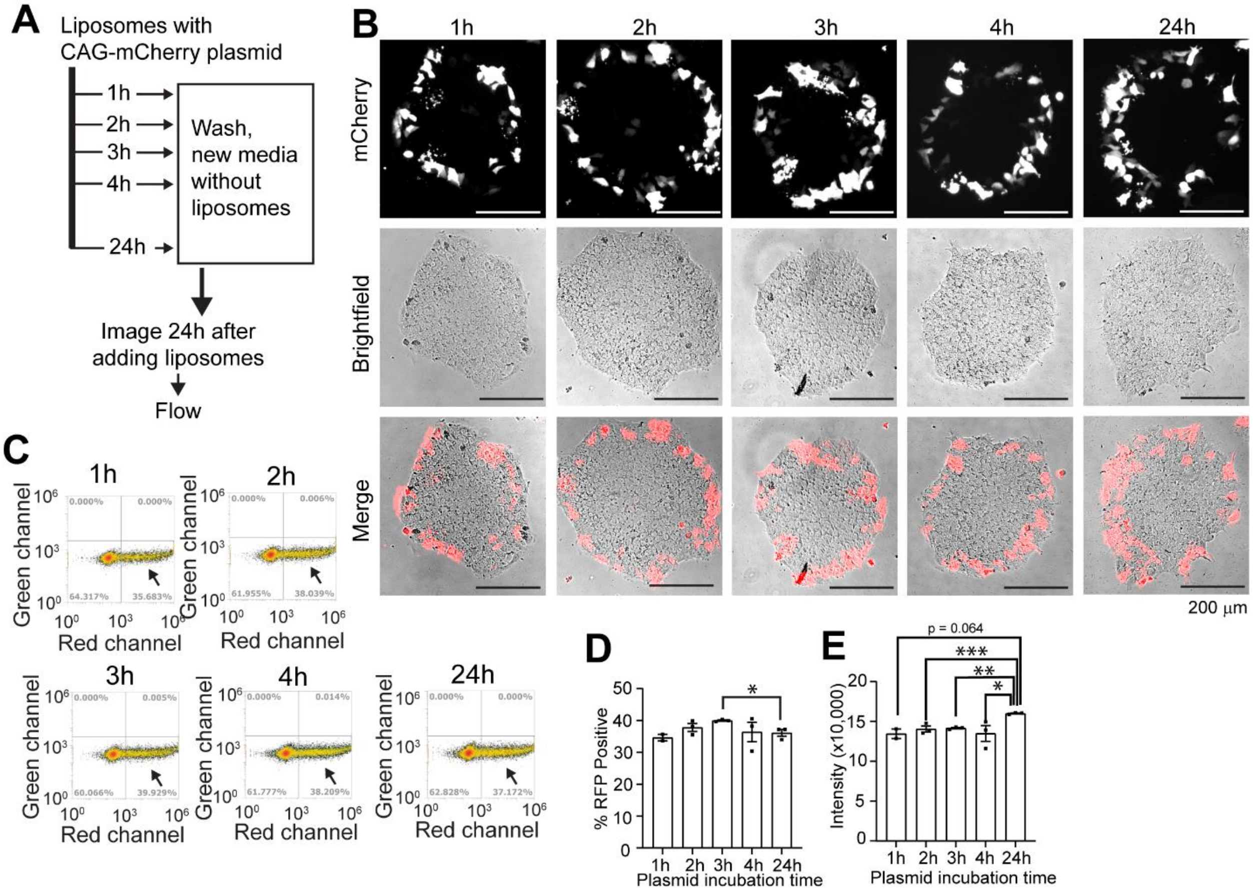

Supplemental Figure S5

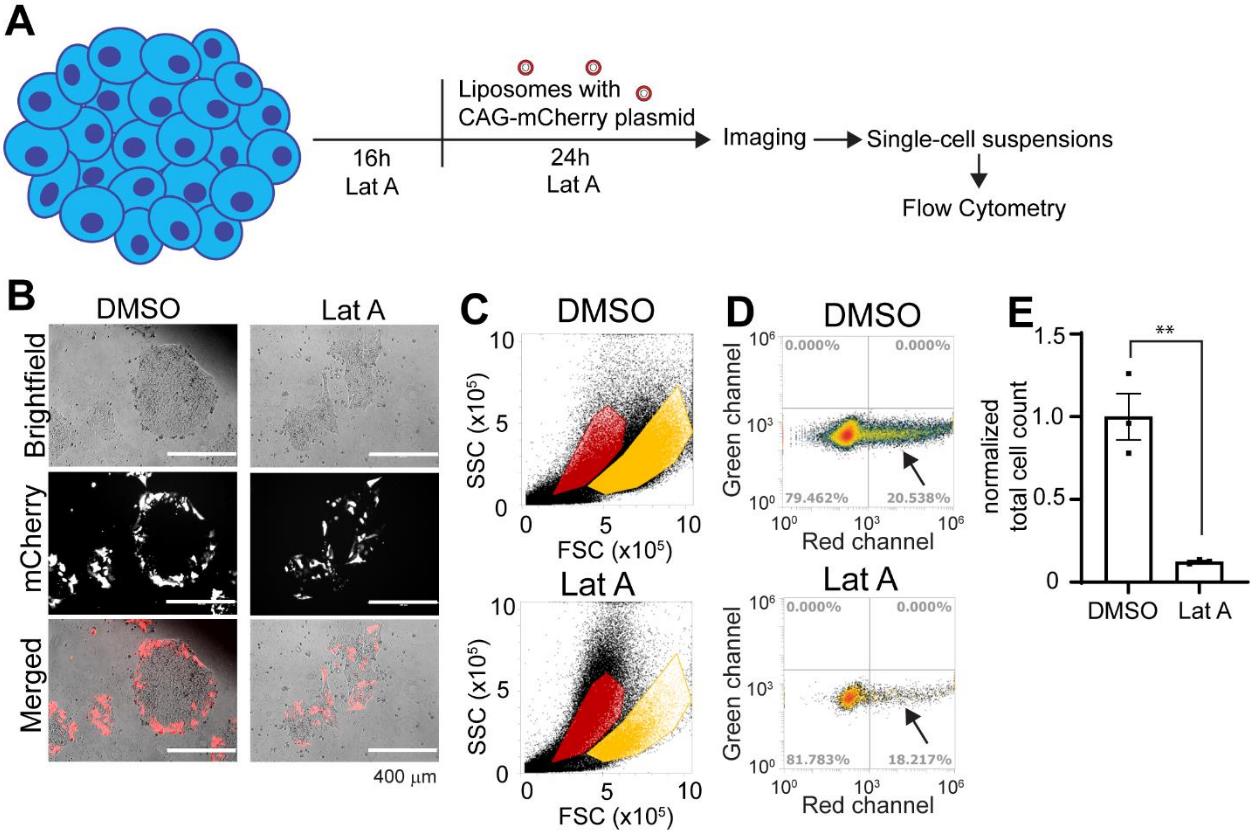

### Supplemental Figure S6

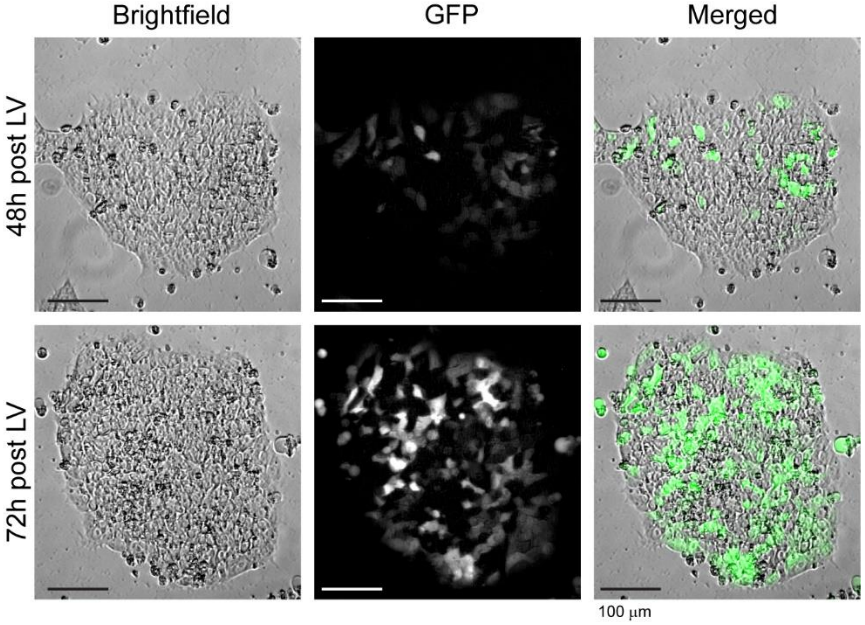
